# Contextual Neural Dynamics During Time Perception in Primate Ventral Premotor Cortex

**DOI:** 10.1101/2024.10.03.616577

**Authors:** Héctor Díaz, Lucas Bayones, Manuel Alvarez, Bernardo Andrade-Ortega, Sebastián Valero, Antonio Zainos, Ranulfo Romo, Román Rossi-Pool

## Abstract

Understanding how time perception adapts to cognitive demands remains a significant challenge. In some contexts, the brain encodes time categorically (as “long” or “short”), while in others, it encodes precise time intervals on a continuous scale. Although the ventral premotor cortex (VPC) is known for its role in complex temporal processes, such as speech, its specific involvement in time estimation remains underexplored. In this study, we investigated how the VPC processes temporal information during a time interval comparison task (TICT) and a time interval categorization task (TCT) in primates. We found a notable heterogeneity in neuronal responses associated with time perception across both tasks. While most neurons responded during time interval presentation, a smaller subset retained this information during the working memory periods. Population-level analysis revealed distinct dynamics between tasks: in TICT, population activity exhibited a linear and parametric relationship with interval duration, whereas in TCT, neuronal activity diverged into two distinct dynamics corresponding to the interval categories. During delay periods, these categorical or parametric representations remained consistent within each task context. This contextual shift underscores the VPC’s adaptive role in interval estimation and highlights how temporal representations are modulated by cognitive demands.

**Significance Statement:** The neural representation of time has long intrigued neuroscientists, particularly how it adapts to cognitive demands. Depending on the task, the brain encodes time either categorically (“long” or “short”) or as precise intervals. While the ventral premotor cortex (VPC) is known for its role in temporal processes, its role in time estimation remains underexplored. Here, we examined how the VPC processes temporal information in primates during a time interval comparison task (TICT) and a categorization task (TCT). The VPC exhibited heterogeneous neuronal responses with distinct dynamics: a linear, parametric relationship in TICT and bifurcated dynamics in TCT. These representations remained consistent during delay periods, underscoring the VPC’s adaptive role in time interval estimation.

## Introduction

Time, an omnipresent dimension of our reality, has intrigued physicists, biologists, neuroscientists, and philosophers for centuries. In physics, time was once considered an absolute constant until Einstein’s theory of relativity (1) unveiled its malleable nature, showing how it can stretch and contract relative to an observer’s velocity and gravitational field. Concurrently, the enigma of how the brain perceives and represents time, remains a formidable unsolved question in neuroscience (2–4).

One of the major challenges in understanding temporal perception is determining how it adapts based on the cognitive demands of a task. In some cases, the brain categorizes time simply as long or short, while in others, it measures the precise duration of an interval on a continuous scale (5, 6). Neural circuits adjust their processing and computation of temporal information depending on the nature of the task (4, 7, 8). While recent research has examined how humans, primates and rodents categorize the duration of stimuli (9–11), there remains significant uncertainty about where and how the perception and memory of time are processed in the brain, and how this information is used for comparison and decision-making (12).

Studies suggest that neurons in the frontal lobe circuits are linked to the perception of time (13–17), with the medial premotor cortex (MPC) playing a key role in tasks related to time and rhythm (4). Additionally, temporally scalable firing patterns have been observed in the prefrontal cortex of rats, suggesting a neural mechanism for interval timing (18). Recently, other brain regions, such as the medial entorhinal cortex in mice, have also been implicated in context-dependent interval timing, pointing to a distributed network for temporal processing (19). However, despite these well-conducted studies on time perception, it remains unclear how neuronal activity encodes a temporal interval marked by two discrete sensory events in the absence of a continuous sensory signal. One proposal suggests that time could be represented through population dynamics, where neural networks capture, retain, and compare temporal information via collective response scaling (14, 15, 20).

Understanding and abstracting physical variables–such as time perception and its influence on cognitive processes–remains a significant challenge, particularly given the prevalence of neurodegenerative conditions that impact these functions (21, 22). Two key theories have emerged to explain how the brain processes time. The first is the dedicated temporal processing hypothesis, which posits the existence of specific circuits functioning as an internal clock, operating through a pacemaker mechanism (4, 23–25). The second is the distributed or intrinsic processing hypothesis, suggesting that temporal processing is an emergent property of local neural dynamics, where cortical region estimates relative time based on the state of neural networks influenced by sensory inputs (11, 26–28).

Building on these theories, and given that monkeys can discriminate between time intervals (29), we sought to determine whether these animals could compare the durations of empty time intervals—intervals marked solely by two discrete pulses without any sensory cues in between. Unlike tasks involving frequency discrimination (30, 31), it has been suggested that an intensity– or integration-based code (32) would not suffice for resolving temporal tasks. For time intervals, both cortical and subcortical circuits must generate a time estimate, filling in gaps between stimuli that mark the boundaries of each interval (33, 34). Once this time estimate is formed (35), it must be stored in working memory for use in comparison and decision-making (15). This raises important questions about the role of less-studied cortical areas in time computation.

In this study, we investigate the role of the ventral premotor cortex (VPC) in processing temporal information. Previous research has identified the VPC as a crucial intermediary between sensory and frontal cortices, receiving projections from sensory regions in the parietal and temporal lobes, as well as from prefrontal and premotor association areas (36–39). During tactile tasks, VPC neurons show distinct and sustained sensory responses (30, 40, 41), and interestingly, they show similar responses to auditory stimuli (42, 43), suggesting that the VPC is capable of processing multimodal inputs and conveying these responses to other premotor regions (44). Supporting this, recent studies in humans suggest that the VPC plays a central role in integrating information and generating abstract representations during speech (45), hinting at its capacity to organize temporal information from different modalities.

To investigate whether the VPC is involved in quantifying time, we recorded neuronal activity from the VPC in two primates during two tasks. In one task, the animal categorized intervals (“long” or “short”), and in the other, compared two-time intervals (*int1* vs. *int2*) and indicated which was longer. We found that in both tasks, VPC neurons are involved in representing and encoding time. In the time interval comparison task (TICT), a linear attractor dynamic emerged, while in the time categorization task (TCT), a U-shaped dynamic allowed the separation of the two categories. Despite sparse persistent activity, VPC neurons contributed to maintaining information during the working memory interval. This suggests that the VPC plays a role in temporal computation, highlighting its capacity to adaptively encode time depending on the cognitive demands of the task at hand.

## Results

We trained two primates to perform two tactile cognitive tasks involving the comparison (TICT) and categorization (TCT) of time intervals. The stimuli were delivered to the skin of one fingertip (Methods). In TICT, the animal compared the duration of a first-time interval (*int1*) against a second-time interval (*int2*), both ranging from 400 to 2000 ms separated by a fixed delay of 2 s (Figure 1A). Following *int2*, a fixed decision maintenance period of 2 s ensued. The animals were then prompted to respond (probe up from the skin, PU) by pressing one of two buttons (PB) to indicate whether they perceived *int1* as longer than *int2* (*int1*>*int2*) or *int2* longer than *int1* (*int1*<*int2*). A typical experimental session consisted in 14 different combinations of *int1* and *int2* distributed across 140 trials randomly presented. Furthermore, the TICT had two variants: an “empty interval”, where the time intervals were marked by brief 20 ms (150 µm) pulses, and a “filled interval”, where a constant oscillatory stimulation (indicate frequency) was maintained throughout the interval. Monkeys were first trained in the filled interval condition (easiest TICT variant) and then switched to the empty time interval (hardest TICT variant). Animals were able to solve the tasks with a performance above 75% in both empty and filled intervals (Figure 1B display performance for only empty time intervals). On the other hand, in the TCT task, the animal received only a single time interval (*int*) (Figure 1C), to be categorized as either “long” or “short”. Similarly to TICT, the presentation of the single *int* was followed by a fixed delay period of 1s until PU, where animals had to indicate the category of the *int* by pressing one of two buttons (PB). In this task, two interleaved contexts were employed: the Short Temporal Range (STR) and the Long Temporal Range (LTR). In the STR context, the animal had to categorize any interval above 300 ms as “long”, with *int* ranging from 50 to 550 ms. In contrast, in the LTR context, any interval above 750 milliseconds had to be categorized as “long”, where *int* ranged from 330 to 1170 ms. Notably, time intervals considered “long” in the STR context would be categorized as “short” in the LTR context. In each TCT session, we presented 12 (or 8) different *int* values across 120 (or 80) trials, respectively, randomly distributed within the STR (or LTR) context. Importantly, the STR and LTR contexts were always presented separately in each training and recording session. The primate successfully learned to categorize intervals correctly with a very high performance in both contexts (Figure 1D). Given the differences in complexity between the two experimental paradigms, we initially trained and recorded neuronal activity from participants during TCT. Subsequently, we trained and recorded them while they performed TICT. To ensure that recordings during both tasks were obtained from the VPC, we acutely inserted recording electrodes through chambers precisely positioned using stereotaxic coordinates derived from prior MRI scans of the monkeys. This allowed us to access identical recording sites in each session.

**Figure 1.**
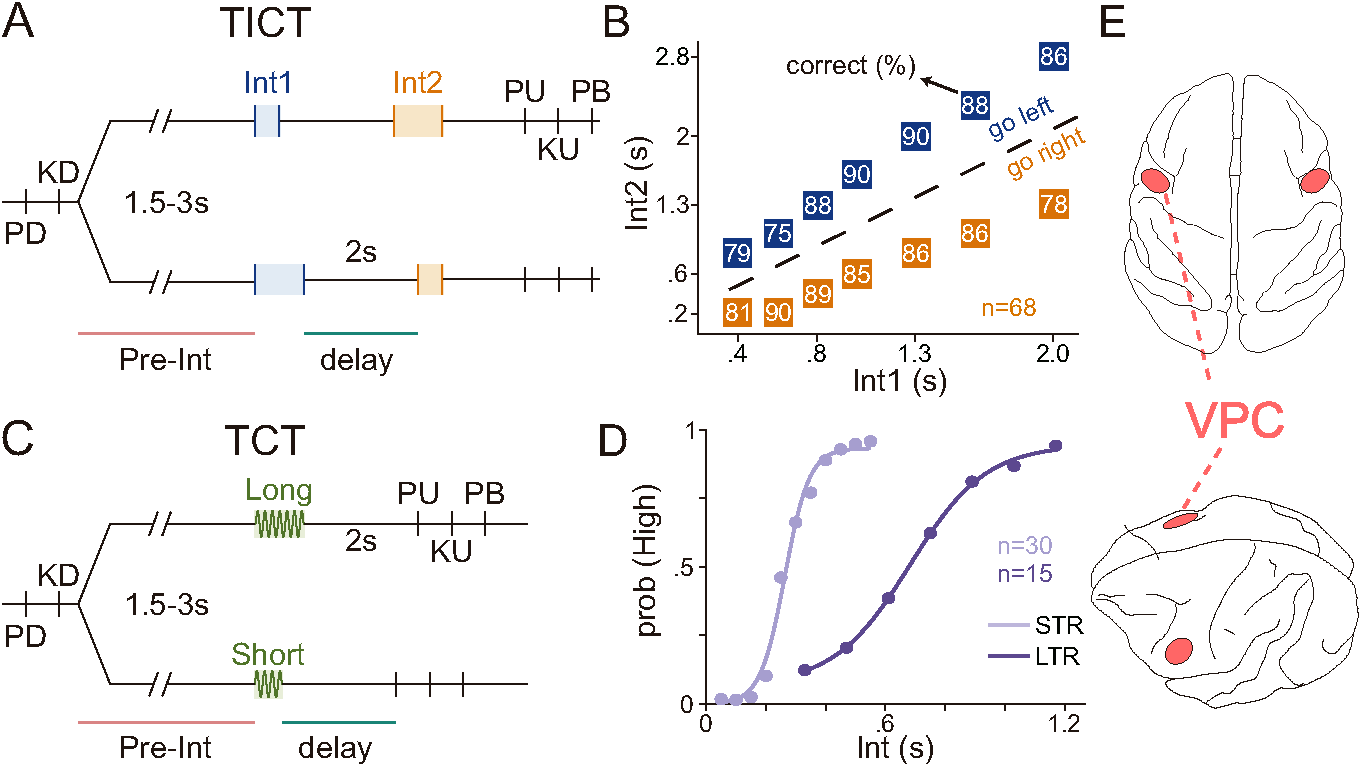
Tasks schematics, performance and recording sites. (A, C) Schematic representations of the Time Interval Comparison Task (TICT) (A) and the Time Categorization Task (TCT) (C). In both tasks, trials begin when a mechanical probe lowers and indents the glabrous skin of a fingertip on the monkey’s restrained right hand (probe down event, “PD”). In response, the monkey places its free left hand on an immovable key (key down event, “KD”). After a variable delay (1.5-3s) following KD, the first interval is presented. (A) In TICT, two pulses mark the beginning and end of each interval. The first interval (*int1*) is followed by a fixed 2-second period, after which the second interval (*int2*) is presented. Following *int2*, another fixed 1-second delay ensues, culminating in the probe up event (“PU”), which serves as the “go” cue for the monkey to release the key (key up event, “KU”). The monkey must then indicate whether *int1*>*int2* or *int1*<*int2* by pressing one of two push-buttons positioned in front of it (push-button event, “PB”). A total of 68 recording sessions were conducted with this task. Correct responses were rewarded with drops of juice. (C) In TCT, the intervals (*int*) to be categorized are indicated by a constant 20 Hz sinusoidal stimulation. After this interval, a fixed 2-second delay follows. Subsequently, the same sequence of events occurs: probe up (“PU”), key up (“KU”), and push-button (“PB”). The animal must indicate whether the interval was “long” or “short”. (B) Psychophysical performance in the TICT. Each box represents a pair of intervals (*int1*, *int2*), with the number indicating the percentage of correct trials for that pair. For each value of *int1*, there are two possible *int2* intervals–one longer and one shorter. The range of intervals extends from 400 ms to 2 seconds. (D) Psychometric curves for the TCT: short temporal range (STR, light violet) and long temporal range (LTR, dark violet). A total of 30 recording sessions were conducted with STR and 15 with LTR. The categorization thresholds are 300 ms for STR and 750 ms for LTR. Notably, intervals classified as “long” in STR are classified as “short” in LTR. (E) Top (upper panel) and lateral (lower panel) views of a monkey’s brain with the recorded areas of the VPC highlighted in pink.

## Neuronal activity during the comparison and categorization of time intervals

In both tasks and for both animals, we recorded neuronal activity from the VPC (Figure 1E), where **n=702** for TICT (empty intervals); **n=132** and **n=47** for STR and LTR in TCT, respectively. Although both filled and empty time intervals were used in the TICT task, our primary focus is on how the VPC encoded time intervals without any sensory reference within the intervals, in other words we explored how this area perceives time in complete abstraction. Thus, the majority of VPC neurons were recorded during empty time intervals in the TICT. Therefore, the following results present data exclusively from the first empty interval (*int1*) during TICT and the unique interval (*int*) during TCT across both contexts. Figures 2, 3, S1, and S2 illustrate the neuronal activity of various VPC neurons in both tasks. Notably, many VPC neurons exhibit dynamics associated with temporal estimation. Each raster panel in the figures is aligned to two reference times: on the left, time zero (0 s) marks the start of *int1* for TICT and the single interval (*int*) for TCT, while on the right, time zero corresponds to the onset of the 2-seconds delay period in both tasks. At the bottom of each raster, the associated firing rate is shown. These activity profiles were obtained by averaging the firing rate from correct individual trials for each condition (*int1* for TICT; *int* for TCT), aligned in the same manner as the rasters above. On the left side, bold colored lines represent the dynamics of each neuron during interval encoding, starting from the onset and continuing to the offset of each interval (colored filled circles). Given that VPC exhibits a marked response latency (31, 40, 41), we extended each dynamic beyond the offsets using shaded colored lines. This extension illustrates, in some cases, the full encoding of the presented empty time intervals by neurons under TICT and the categorical splitting by neurons under TCT. Conversely, the activity profiles on the right side represent the neurons’ significant encoding of time intervals during the delay period in TICT or the decision-delay period in TCT. In these profiles, orange markers indicate time bins where neurons carry significant information about the time intervals, while dark green markers denote time bins where significant encoding of the time intervals occurs.

**Figure 2.**
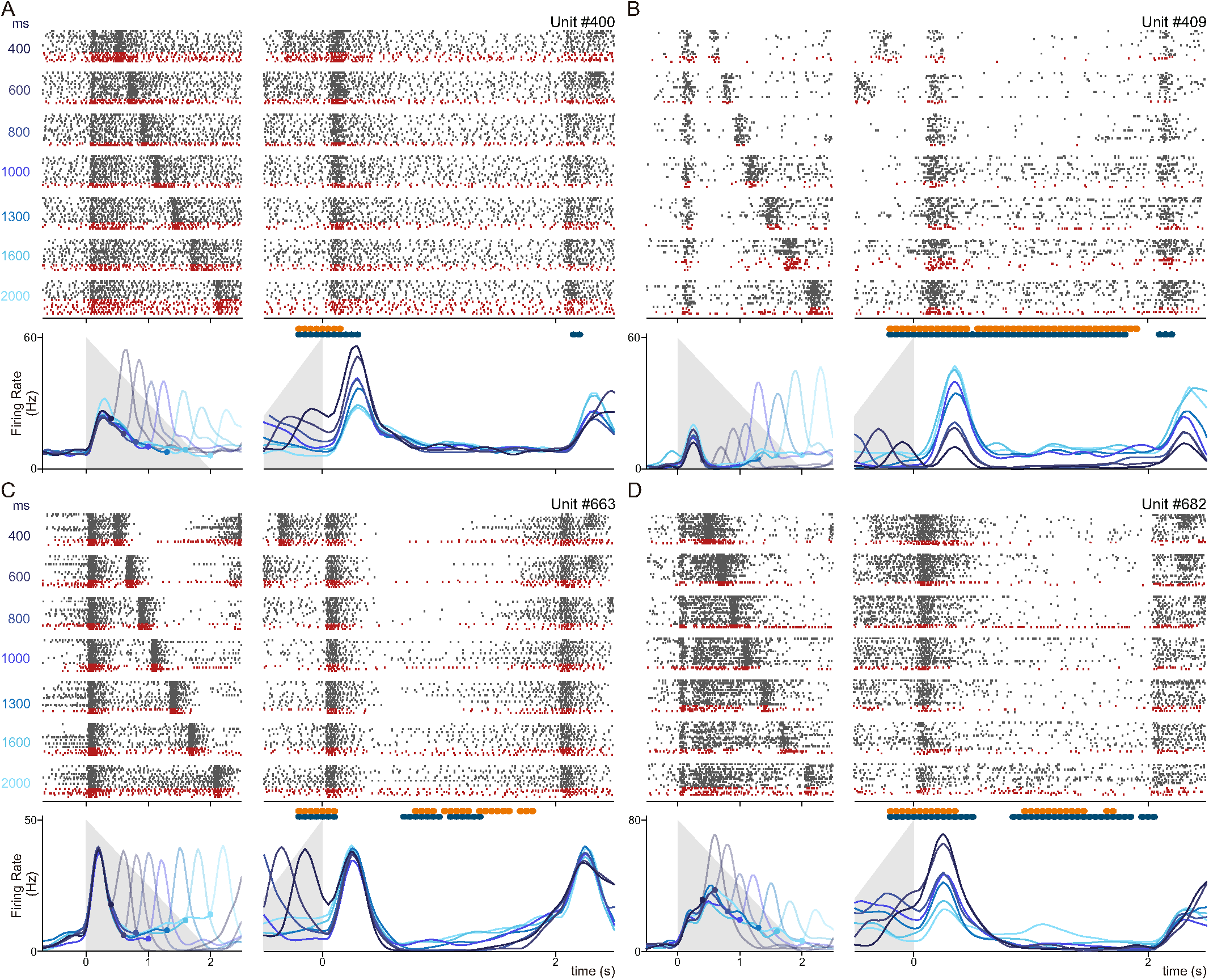
Neuronal activity profiles from the VPC during TICT. (A-D) Raster plots and corresponding firing rate profiles of four individual neurons recorded in the VPC during TICT. In the raster plots, black and red ticks represent spikes during trials with correct and incorrect answers, respectively. For each neuron, the left raster plots references activity aligned to the beginning of the first interval (*int1*, t=0s), while the right raster plots references activity aligned to the beginning of the fixed 2-second interval period. Trials are grouped by the duration of *int1*, ranging from 400 to 2000 ms. Firing rate plots were computed as the mean of correct-answer trials for each *int1*. Grey triangles denote the increasing values of *int1*, illustrating how the firing rate varies with changes in interval duration during successful trials. Orange and green markers above the activity profiles indicate time windows with significant (*p<0.05*) mutual information and F-test (coding) related to *int1*, respectively, highlighting moments where the neuron’s firing pattern shows a statistically significant correlation with the duration of *int1*. Notably, neurons A, B, and D encode the identity of the interval at the onset of the memory interval, exhibiting a parametric response in their firing rates associated with *int1*. Additionally, neurons B, C, and D carry temporal information at various moments throughout the memory period.

We observed that many neurons modulated their activity during the task components of TICT (Figures 2A-D and S1A-D). If we examine the activity profile (left subpanels) of some neurons, such as those in Figure 2C and S1C, we can observe a strong peak of activity at the onset of the time interval and another equally marked peak at the end of each time interval when considering VPC’s response delay (colored shaded lines). In this case, both neurons are strongly coding the beginning and end of each time interval, as shown in their raster plots above. However, at the beginning of the delay period (right subpanels), both neurons lack modulation for a particular condition. However, this changes over time for the unit in Fig. 2C, where a marked modulation for long intervals is exhibited during most of the delay, then switching its preference to short intervals at the end of this epoch. Other neurons—illustrated in Figures 2A, 2B, 2D, S1A, S1B, S1D— also modulated their activity at the end of *int1*, with their firing rate correlated with the interval’s duration. Differently to units in Figs. 2C and S1C, here we identified neurons with positive coding, where longer intervals triggered higher firing rates (Figures 2B and S1A), and neurons with negative coding, showing higher firing rates for shorter intervals (Figures 2A, 2D, S1B and S1D). This might suggest that the activity of these neurons at the end of the first interval can be used to decode the proper length of the time interval. Importantly, some neurons–such as those depicted in Figures S1A and S1D– showed complex dynamics associated with time estimation that are not easily discernible through simple visual analysis of their activity. Therefore, these results focusing on neural activity elicited from empty interval presentation (considering VPC’s latency) indicate that neurons from this area play a significant role in the encoding of time intervals. When translating this analysis to the memory period, the epoch between the end of *int1* and the beginning of int2, it is noticeable how some of these neurons continued to encode time information during the early part of this delay (Fig. 2A, B & D and S1B). Thus, while the primary responses of these neurons appear to encode *int1* during the final part of the interval, they also retain part of the information at the onset of the working memory period of the TICT, arguably for comparison purposes with *int2*. Notably, although they were not the majority, some units depicted in Figs. 2B & C maintained *int1* information throughout most of the memory epoch, exhibiting a parametric encoding of the presented time intervals.

When comparing the neuronal responses of VPC during TICT with those during the TCT, it becomes evident that there are similar patterns of negative and positive encoding of time intervals, as observed during STR (Figure 3A & B) and also in Figure S1 (panels F and H, respectively). Nevertheless, during stimulus presentation the activity response profiles of these neurons also showed a clear encoding for the interval category (shaded lines). Some units showed higher responses for low categories (Figures 3A and S1F), while others perform similarly but for high categories (Figures 3B and S1H). Conversely to the response profiles during TICT, VPC neurons during TCT (STR and LTR) exhibited similar activity values for intervals within each category (short or long) and reflect the encoding of that specific category during the decision memory period. Thus, while time is significantly encoded in VPC neurons in both tasks, the encoding is parametric in TICT and categorical in TCT. Among the neuronal responses described above, other neurons (Figures 3C, 3D, S1E, S1G) showed more complex dynamics, presumably involved in encoding time interval estimation. Similarly, these encoding patterns were also observed when presenting LRT (Figure S2). Several neurons from this paradigm exhibited categorical encoding during time interval presentation, continuing in the decision memory delay (Figure S2B, S2C and S2E).

**Figure 3.**
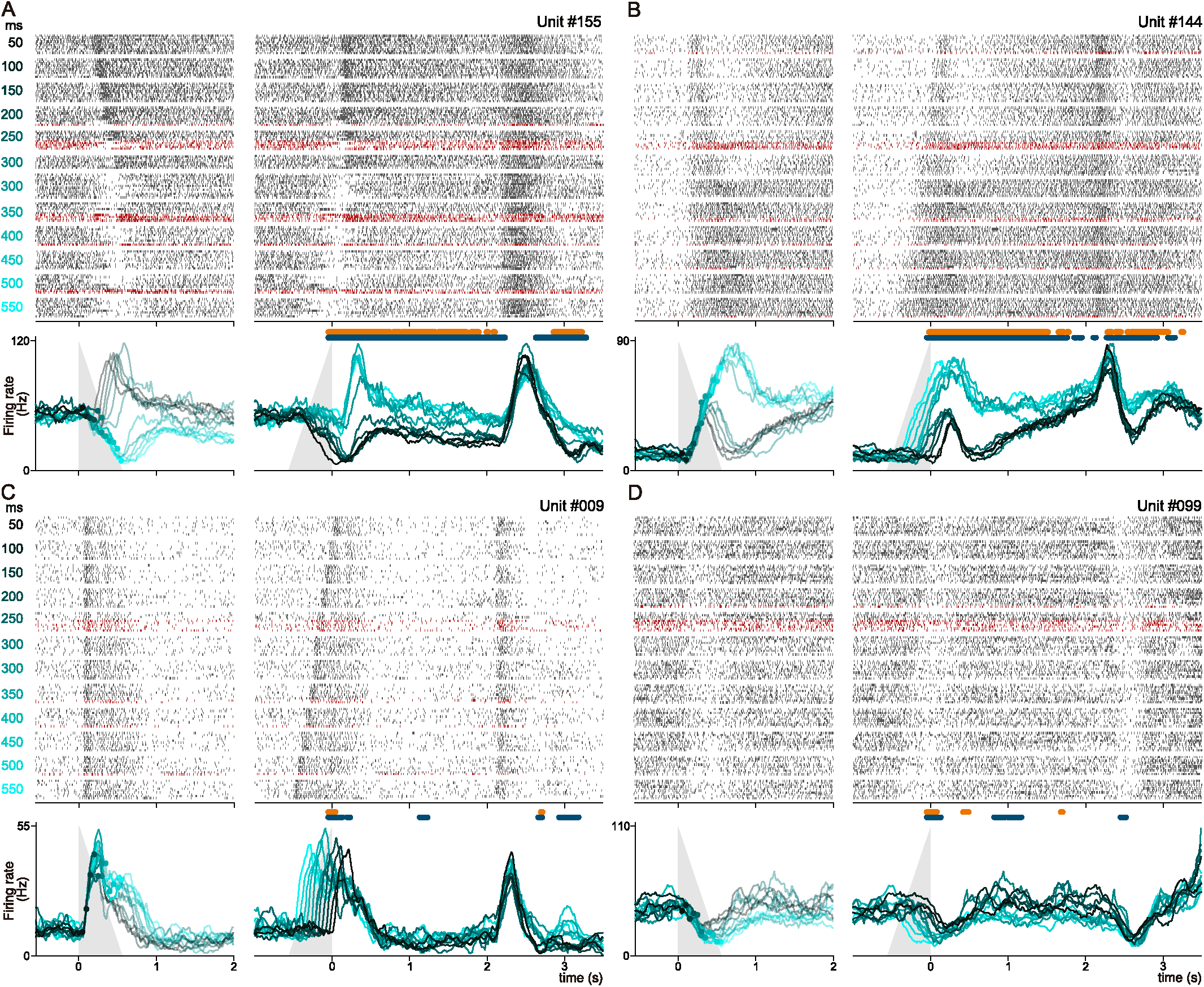
Single neurons’ activity from the VPC during SRT of TCT. (A-D) Raster plots and corresponding firing rate profiles of four individual neurons recorded during SRT. In the raster plots, black and red tick marks represent spikes during trials with correct and incorrect responses, respectively. For each neuron, the left raster plots references activity aligned at the beginning of the unique interval (*int*, t=0), while the right raster plots aligns to the start of the fixed 2-second decision interval. Trials are labeled by the duration of *int*, ranging from 50 to 550ms. In this set, all intervals over 300 ms should be categorized as “long”. The 300 ms trials (threshold interval) were divided based on the animal’s decision. Firing rate plots we computed as the mean among correct-answer trials for each *int* used. Grey triangles denote the crescent *int* values, illustrating how the firing rate changes with variations in interval duration among successful trials. The orange and green markers on top of neuronal activity represent time-windows with significant (p<0.05) mutual information and F-test (coding) related to *int*, respectively. Neurons A and B encode the category of the interval, particularly during the first part of the decision delay, while neurons C and D display dynamics more involved in quantifying the duration of the intervals, as seen in the raster plots.

Despite the high heterogeneity of neuronal responses across tasks and conditions, our results indicate that VPC neurons play a role in encoding time intervals in both the TICT and TCT. Importantly, the encoding patterns differ significantly depending on the task context. In TCT, neurons not only encode the interval but also sustain categorical information throughout the memory-decision period. In contrast, during TICT, VPC neurons primarily encode the interval (*int1*) and mostly maintain this information only in the early phase of the working memory period (between *int1* and *int2*). This distinction suggests that while VPC neurons consistently contribute to encode time, their activity profiles adapt based on the cognitive demand of the task (compare or categorize).

## Neuronal encoding during comparison and categorization of time intervals

We further investigated how neuronal populations code time interval during TICT and TCT (Figure 4). By analyzing the collective activity, we identified distinct patterns that reveal how this area processes temporal information in different contexts. Focusing on the first interval of TICT, we organized the neurons based on the temporal interval during which each exhibited its highest activity. This arrangement produced a graph similar to the one shown in Figure 4A (left). To compare neurons with different response ranges, we normalized each neuron’s activity. Remarkably, we observed a continuum of responses: the neuronal population of VPC exhibited peak activity across the entire range of the presented time intervals. In the middle panels, we display representative normalized responses of neurons with peak activity at the beginning (#1), middle (#261), and end (#058) of *int1*. These results suggest that different neurons in the VPC are tuned to specific time points within the interval, with each neuron’s peak activity marking distinct time points. For example, if neuron #058 does not fire during a trial, it signals to the circuit that the elapsed time is less than 1.5 seconds. A similar dynamic was observed during TCT for both STR (Figure 4B left) and LTR (Figure S3 left) contexts. Once again, these panels include representative neurons from STR and LTR showing normalized peak activity at different times points during the interval. These findings suggests that the VPC generates dynamic, time-evolving activity patterns that code temporal information in both tasks.

**Figure 4.**
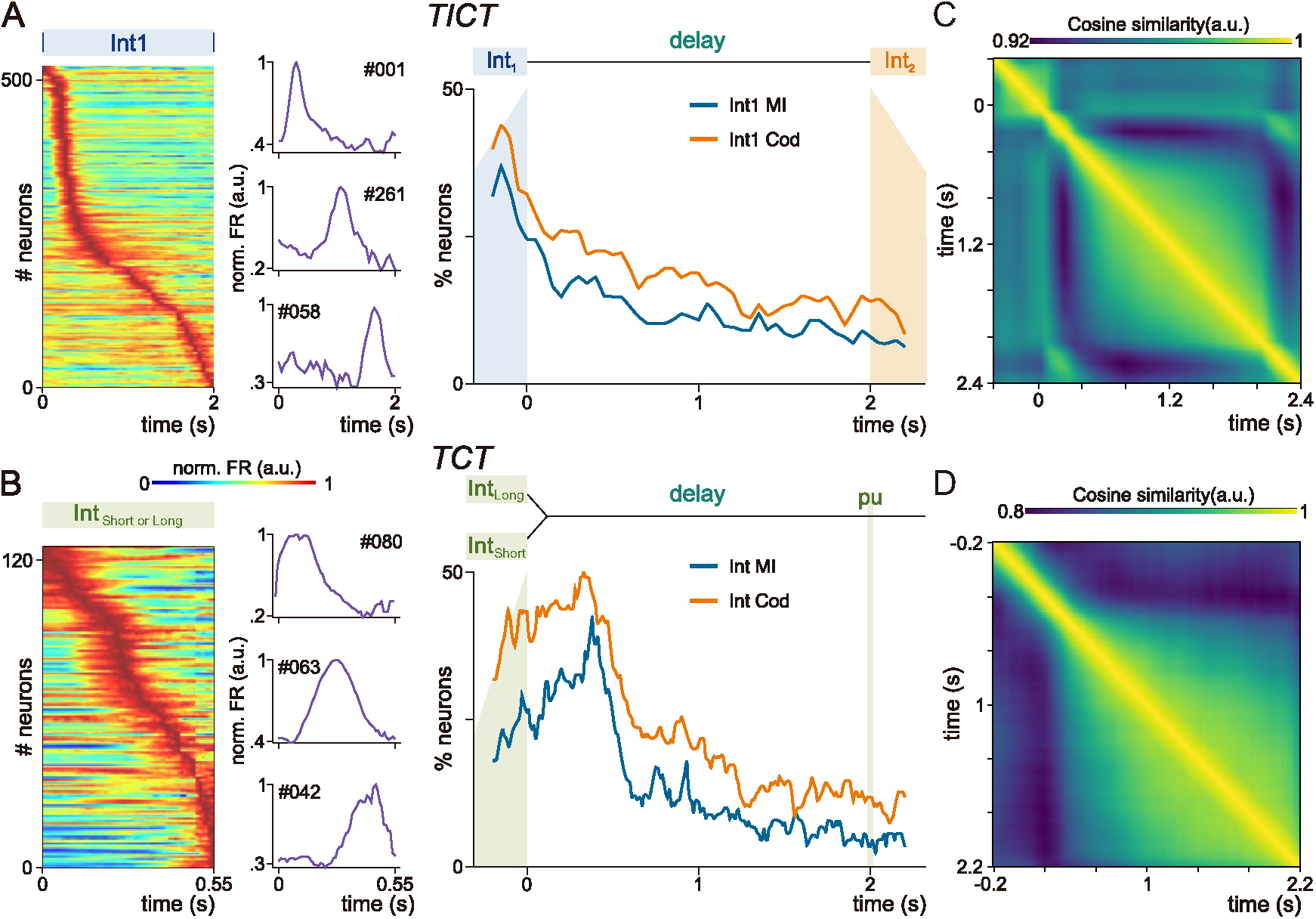
Estimation and maintainance of temporal information by the population of VPC neurons. (A, B) **Left**: Dynamics of individual neurons during the presentation of the interval to be compared in TICT (A) or to be categorized in TCT (B). Neurons with more than 5σ from their basal activity were normalized and positioned on the color-coded graph by the time of their peak activity. Neurons at the top respond earlier, while those at the bottom respond later. In both cases, a “moving bump” dynamic is used by the population of neurons as a strategy for decoding time intervals. **Middle**: Example of neurons with normalized peak activity at three different specific times within the range of time intervals. **Right**: Percentage of neurons with significant encoding during the working memory delay (A, TICT) and decision maintenance delay (B, TCT). Mutual information (blue) and F-test (orange) were used to identify neurons with significant time interval encoding (p<0.05). In TICT, after the interval presentation, the animal must compare *int1* vs *int2*. In TCT, following the interval and after PU, the animal is prompted to report its decision. In both tasks, a larger number of neurons exhibit significant encoding at the start of the delay with this number gradually decreasing over the 2-second period. Nevertheless, a substantial level of significant encoding persists throughout the delay in both tasks. (C, D) Cosine similarity matrices of population states in TICT and TCT. Comparison of population states between two distinct times, t1 and t2, during the working memory delays in the TICT (A) and the decision maintenance delay in the TCT (B). At each time point, a population vector is constructed, with each line representing a neuron’s activity across different intervals (*int1* or *int*). The cosine of the angle between vectors from two distinct times is plotted, with yellow indicating high similarity and blue indicating low similarity among population states. Importantly, in both tasks, there is a clear separation between the population state at the end of the interval and the beginning of the delay, with dynamics during the subsequent maintenance periods.

We next investigated whether VPC neurons coded temporal information during the delay period of both tasks. Using mutual information and analysis of variance (ANOVA), we calculated the percentage of neurons that significantly carried information associated with the interval duration in TICT and TCT during the two-second delay. In TICT (Figure 4A, right), most neurons encoded interval information at the beginning of the delay period, and a considerable number continue to do so throughout the entire two-second retention period. Maintaining this information is crucial in TICT, as the animal needs to compare the *int2* with *int1*. In the following sections, we will present evidence demonstrating that this information is maintained with a parametric, approximately linear code within the neuronal population. Similarly, during TCT, where the animal must retain the interval category before pressing the correct button, a high percentage of neurons encoded the interval category at the beginning of the delay in both the STR (Figure 4B, right) and LTR (Figure S3, right) contexts. However, the number of neurons involved decreases during the decision maintenance period. These findings so far suggest that, in both tasks, VPC neurons are involved in encoding temporal information during memory delays, particularly at their onset. It is important to emphasize that although neurons significantly encode time information in both tasks, the coding strategy differs between them. As we will demonstrate in the following sections, a parametric code emerges at the population level in TICT, whereas a categorical population code is observed in TCT.

## Sensory versus Memory Population Responses in Ventral Premotor Cortex

To investigate how intervals are coded during the sensory versus memory periods, we compared the neural population states across different time points (46–48). We constructed N-dimensional vectors representing population activity for each interval, where each element corresponds to the activity of a neuron in a particular class. Using cosine similarity (*CS*), we compared these vectors at various times to quantify the similarity in population representations between different periods (see Methods). This analysis results in time-by-time (*TxT*) matrices, enabling us to systematically measure the dynamics of these effects.

In Figure 4C, we present this matrix for the working memory period of TICT. Similarly, in Figures 4D and S3B, we compute these matrices for TCT in both the STR and LRT contexts. This analytical approach allows for a direct comparison of population dynamics across distinct tasks. Since the working memory intervals in TICT and the decision memory period in TCT both last 2 seconds, these matrices can be directly compared. Across scenarios, we observed a distinct separation at the population level between the period spanning from the end of the interval to the onset of the delay, as compared to the rest of the delay. Notably, during the final period of the delay, the population dynamics remained consistent, yet distinctly different from those observed during initial part of the delay. This finding illustrates a clear decoupling of dynamics between the sensory and memory periods in both tasks.

It is important to mention that the separation between early and persistent neuronal activity during memory periods is not novel (41, 49, 50). However, the context of temporal coding provides a unique framework for understanding these dynamics, distinguishing our findings from previous studies.

## Contextual Time Interval Representation in VPC Neuronal Populations

Throughout this study, we have delved into the complex role of the VPC in encoding time intervals, emphasizing the challenges posed by the heterogeneity of neural responses observed in both categorization and comparison tasks. Despite this diversity, our analyses suggest that some underlying mechanisms in the VPC may be shared across different contexts, while others vary depending on the specific task demands. In this section, we focus on studying these latent dynamics to draw comparisons across different contexts. To achieve this, we applied principal component analysis (PCA) to the neuronal activity recorded from the VPC during stimulus presentation in both tasks (see Methods). This approach revealed significant principal components (PCs) that illuminate how interval information is encoded and represented (51). In Figure 5, we present these significant PCs calculated during *int1* of TICT (Figure 5A) and during the single interval (*int*) of TCT (Figure 5B). Importantly, only the activity from the intervals and the subsequent 200 ms—accounting for the VPC’s response latency (31, 40–42) —was used to construct the covariance matrices from which the PCs were derived. We chose to calculate the interval PCs, extending 200 ms into the delay, to ensure that the response latency observed in previous tasks was adequately captured. Reducing this period would exclude the second pulse or the last part of the interval from the computation, potentially omitting crucial neural activity involved in the time interval estimation.

**Figure 5.**
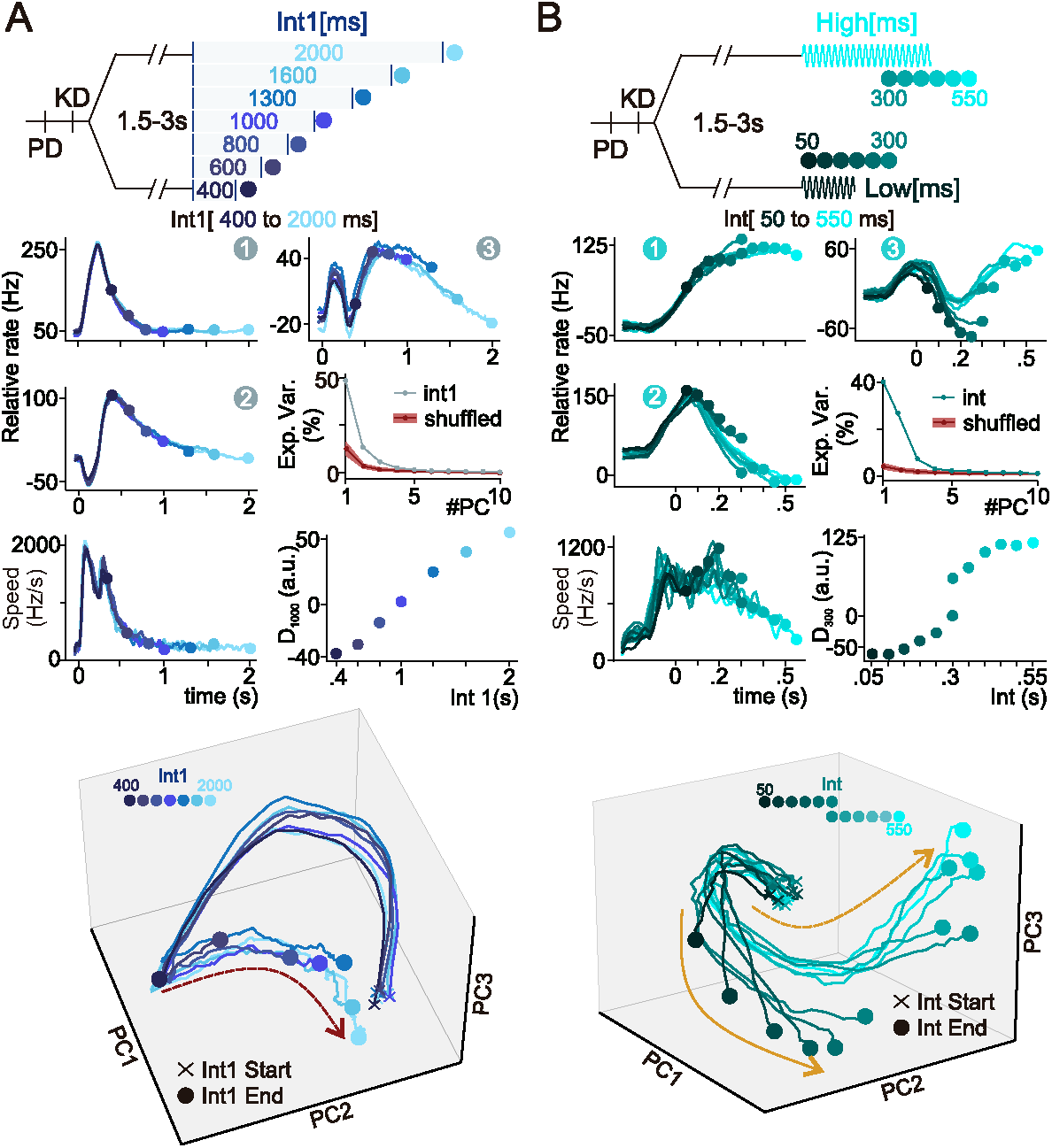
VPC contextual population dynamics in time interval estimation periods. (A, B) VPC’s population dynamics during interval presentation in TICT (*int1*) and TCT (*int*). Principal components (PCs) were derived from applying principal component analysis (PCA) to the covariance matrices constructed by using only the activity of each interval presented plus the first 200ms of the memory delay (VPC’s response delay) and ordered according to their explained variance (EV) for both, TICT (PC1: 49.6%, PC2: 14.4%, PC3: 5.3%) and STR (PC1: 40.2%, PC2: 24.3%, PC3: 7.1%) contexts. **Top:** Schematic representations of the TICT (A) and TCT (B) during the presentation of *int1* and *int* (STR), respectively. **Middle:** Individual projections of neural activity framed in each interval (plus 200 ms) onto the first three significant PCs for both tasks. **Bottom:** 3D-phase diagram using the projections from the three significant dimensions shows the evolution of the dynamics during *int1* in TICT (A) and *int* in STR (B) in the state space. In TICT, dynamics commence (crosses) in the same region and end (circles) in different positions of the state space depending on *int1* duration. Similarly, in the STR context, the dynamics begin closely together and progressively diverge based on the interval duration, while also segregating according to their respective categories. The panels above the phase diagram show the evolution speed of the neuronal population, revealing that the circuit initially evolves rapidly in both tasks before slowing down, suggesting an early rapid adaptation followed by stabilization. Additionally, the distance of the population state at the end of the 1000 ms interval (D_1000_) in TICT and the 300 ms interval (D_300_) in TCT is plotted, highlighting key differences in neuronal representation. In TICT, the interval is represented parametrically and approximately linearly, indicating continuous encoding based on duration. In contrast, in TCT, the intervals are distinctly separated by category, reflecting a context-dependent shift in encoding.

In Figure 5A, we present three significant principal components (PCs) for the TICT during the *int1* presentation. To determine their significance, we used permuted matrices (see Methods), which revealed that only these three components were statistically significant. Notably, the final positions along these PCs allow us to decode the interval duration. To further illustrate this, we plotted the evolution of the population dynamics in a 3D phase diagram (bottom panel). All trials begin from a common starting point (crosses) and conclude at different positions depending on the interval duration (filled circles). This representation shows an approximately parametric coding pattern during the final period, enabling the interval to be decoded based on the population’s final position.

To further analyze this parametric representation, we constructed a vector representing the signed distance between the population’s final positions in *int1* and the reference vector (see Methods) (15). Specifically, we used the final position of the 1000 ms interval as the reference to compute the signed distance D_1000_. The results of this computation are displayed in the D_1000_ vs. *int1* panel. Interestingly, this projection exhibits an approximately linear parametric relationship, demonstrating a systematic pattern in how the population codes time. Additionally, the speed of the population dynamics (as detailed in the Methods section) is initially high at the beginning of the interval and gradually decreases as the interval progresses, reflecting the evolving neural activity during the task.

When we repeated these computations for the SRT within TCT, focusing on the interval presentation period (Figure 5B), we obtained a similar number of significant PCs as in TICT. Although the duration of intervals in this task is shorter than those in TICT, the dynamics observed in the three PCs during the first 500 ms are comparable between both tasks. However, when we generated the 3D-phase diagram (bottom panel), a significant change emerged. As before, all dynamics start from the same point (crosses) but end in different regions of the diagram (circles), depending on the category to which the interval belongs. Importantly, in the graphs, the 300 ms category appears twice because, in some trials, the 300 ms interval was categorized as “short,” while in others, it was categorized as “long”. This variability also provides an implicit and rich opportunity to study error processing within the circuit, as the same sensory input leads to different behavioral outputs. Despite the interval duration being identical, the final positions in the phase diagram differ significantly based on the categorization, indicating that the VPC circuit differentiates the dynamics associated with each interval according to the perceived category.

To explore this phenomenon further, we generated a new vector to project the activity at the end of the interval (*int*) in the TCT. Specifically, the reference vector was constructed using the neural representations of 300 ms intervals categorized as “short” at the end of the interval. We then computed the distance between this reference vector and those obtained from each interval (*int*) (see Methods). The projection of the population activity at the end of the interval is shown in the D_300_ vs. *int* panel, revealing a clear separation between points categorized as “short” and “long.” Consistent with the comparison task, the speed of the population dynamics is high at the beginning of the interval and decreases as the interval progresses.

When repeating these calculations for the LTR in TCT, we also obtained three significant PCs. Despite the smaller number of neurons and increased variability in dynamics, the 3D phase diagram revealed an even clearer separation among the different categories. Notably, the 750 ms trials, which serve as the categorization threshold in this context, exhibited a pronounced divergence in population activity depending on the perceived category. As before, we generated a reference vector using these trials, and the projection of the final dynamics (D_750_ vs. Int) showed an even clearer separation than in the STR.

These findings represent a highly significant outcome of our work, highlighting the adaptive dynamics of the VPC in processing temporal information. Although the VPC is involved in both comparing and categorizing intervals across different ranges, the geometric representation it employs during each task changes drastically depending on the context and cognitive demands. In the comparison task, a linear and parametric coding scheme emerges at the end of the interval, reflecting a direct relationship between the population state and the perceived time duration. In contrast, during categorization, the dynamics diverge significantly based on the temporal threshold or reference used, resulting in distinct geometric patterns. Thus, in the comparison task (TICT), the final population state is associated with the perceived interval duration, whereas in the categorization task (TCT), the final position corresponds to the perceived category.

## Context-Dependent Working Memory Representation in the VPC Neuronal Populations

Building upon our previous findings, which demonstrate how the VPC network flexibly adjusts its processing to encode time intervals either parametrically or categorically, we now turn our attention to how this information is biologically and geometrically maintained during the delay period—a phase whose role varies significantly depending on the task. In TICT, maintaining the representation of *int1* is critical for comparing it with the subsequent interval (*int2*). Conversely, in TCT, preserving the representation of the category is essential for decision-making. Our observations from Figures 3 and S3 revealed a reduction in the number of neurons that retain information as the delay progresses. This finding raises a crucial question: Does a consistent population representation exist in the VPC during the working memory phase across both tasks?

We computed the PCs by generating a covariance matrix from the neuronal activity during the delay period, enabling us to closely examine how the neuronal population maintains information about *int1* in the TICT and the single interval (*int*) in the TCT (see Methods). Analyzing the dynamics during TICT (Figure 6A), we found that only three PCs were significant throughout the memory period. Notably, the third component exhibited a continuous gradient corresponding to the specific interval duration that needed to be maintained. This became more apparent when we plotted the evolution of the population dynamics in a 3D phase diagram (Figure 6A, bottom). The trajectories illustrate the beginning of the delay period (circles), aligning with the end of interval encoding observed in Figure 5A. Remarkably, the parametric representation observed during the interval persists throughout the working memory period, as it is also reflected in the evolution of the speed and distance between trajectories. This is evidenced by the consistent parametric representation at the end of the delay (squares), mirroring the representation seen at the onset. These results suggest a robust mechanism by which the VPC maintains temporal information over time.

**Figure 6.**
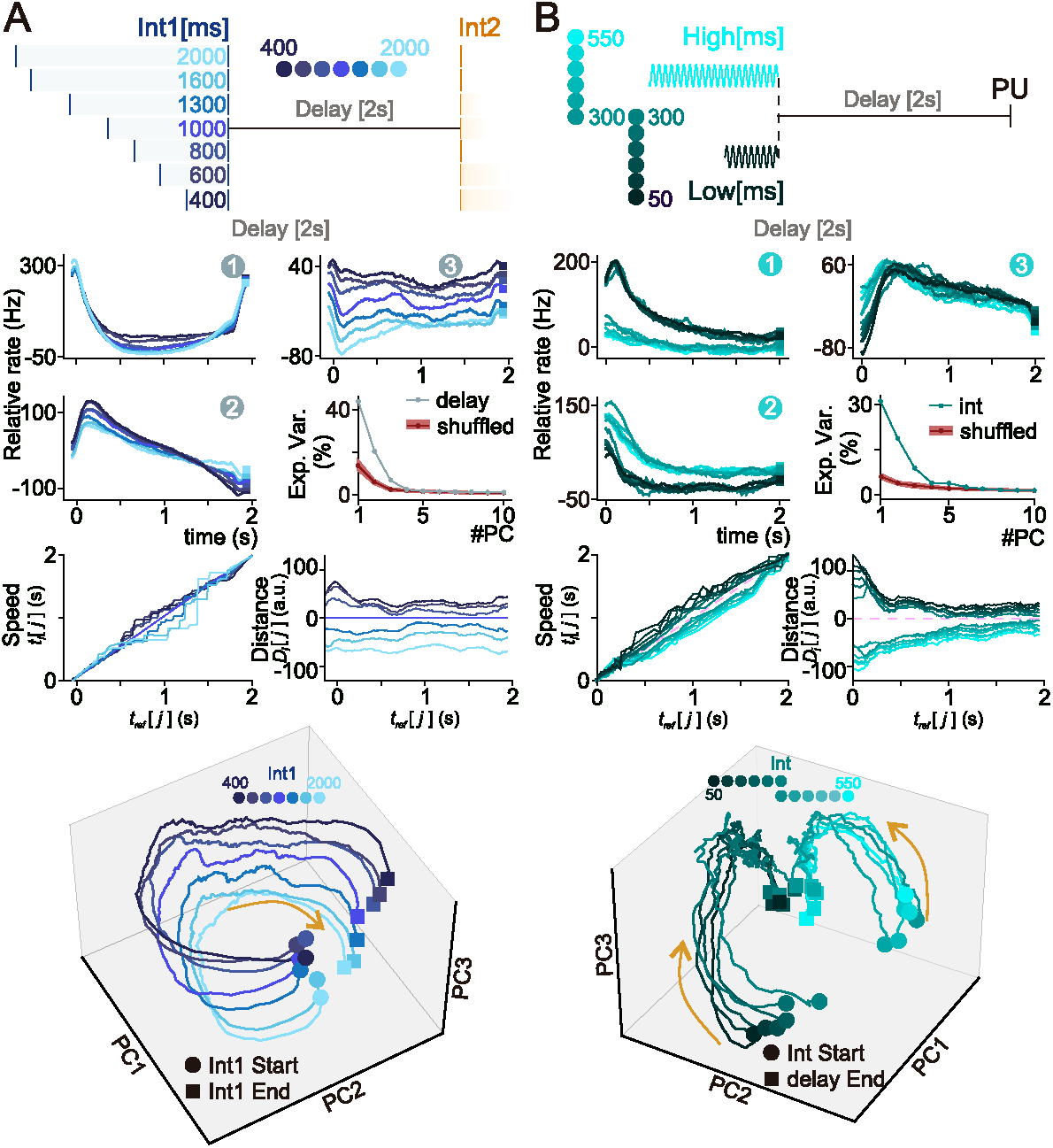
Context-dependent dynamics in VPC during time interval information retention periods. (A, B) VPC’s population dynamics during working memory (TICT) and decision maintenance (TCT) delays. PCs were derived from applying PCA to the covariance matrices constructed solely from activity recorded during the 2-second delay periods (shifted 200 ms forward in the timeline) and ordered according to their explained variance (EV) for both, TICT (PC1: 44.1%, PC2: 20.6%, PC3: 4.8%) and STR (PC1: 31.1%, PC2: 18.7%, PC3: 8.8%) contexts. **Top:** Schematics for TICT (A) and STR (B) during their respective 2-second delay periods. **Middle:** Individual projections of neural activity framed in each task dealy onto the first three significant PCs. Notably in TICT, the dynamics followed a parametric trajectory in PC3, while in STR, PC1 and PC2 exhibited categorical separations. **Bottom:** 3D phase diagram using the projections from the three significant dimensions shows the evolution of the dynamics during each task delay. Beginning and end of the delay are marked by circles and squares, respectively. Note that the dynamics begin at different positions, determined by the endpoint of the interval period (circles in Figure 5). Speed panels above the phase diagram showcast the evolution of the neuronal population, which evolves constantly and independently of the interval represented. Trajectories distance from the 1000 ms (TICT) and the 300 ms (STR) projection vectors, revealed a parametric representation throughout the entire delay for TICT and a two-category representation for STR. This demonstrates a contextual differentiation between tasks in the delay, which relies on the information to be maintained.

We extended our analysis to examine the neuronal activity during the decision maintenance delay in the SRT (Figure 6B). In this case, the first two PCs revealed a clear and persistent separation of the dynamics into two distinct categories, which continued throughout the delay period and only began to converge as the execution period commenced. The phase diagram at the bottom of Figure 6B illustrates that the dynamics initially start divided based on the interval category and maintain this division throughout the delay. This separation, already evident at the end of the interval as depicted in Figure 5B, is consistently preserved within the VPC population during the delay. The panel displaying the distance between trajectories over time shows that the two categories remain distinctly separate throughout the entire delay period. Moreover, the evolution speed of the dynamics is consistent across both categories, indicating a uniform dynamic within the VPC regardless of categorization.

Additionally, when applying the same analytical approach to the LTR (Figure S4B), similar findings emerged: the dynamics consistently maintained a separation between the two categories throughout the delay period. This consistent separation across both STR and LTR contexts highlights the VPC’s robust capability to handle categorical distinctions in temporal tasks, maintaining clear and distinct neuronal representations across varying temporal scales.

Therefore, we demonstrate that the VPC not only represents temporal information in a context-dependent manner during stimulus presentation but also maintains this representation throughout the delay period. In TICT, the VPC preserves information parametrically in memory, with neural dynamics evolving in an orderly fashion corresponding to the specific interval durations. Conversely, in TCT, the VPC’s dynamics distinctly separate the intervals into two clear categories, maintaining this categorical separation throughout the decision delay period. This adaptive ability to manage different types of temporal information underscores the VPC’s crucial role in both encoding and sustaining memory processes.

## Discussion

We have provided compelling evidence that the VPC plays a crucial role in coding time intervals and maintaining interval-related information in memory. Through recordings during the TICT and TCT, we observed that individual neuron activity in the VPC exhibits a high degree of response heterogeneity, similar to patterns observed in other frontal areas during various cognitive tasks (11, 14, 46, 52). Interestingly, in both tasks, the population representation during interval estimation appears to be decoupled from the maintenance periods, with the two processes exhibiting decorrelated dynamics. By applying PCA, we condensed the population’s diverse dynamics into a few meaningful latent signals. This procedure revealed that the dynamics of VPC shifts significantly depending on the task context. <u>For the avid reader:</u> it is important to clarify that here, we use “context” to refer to the specific conditions under which the interval-related decision can be formed. In TCT, a categorical judgment can be reached immediately after a single interval is presented, fostering a categorical representation. In contrast, during TICT, no decision is possible until the second interval is presented, necessitating a linear code that must be maintained until further temporal information becomes available. As we have shown, in TICT, a linear and parametric population code emerges at the end of the interval and the onset of the memory period, reflecting a direct relationship between neuronal activity and the perceived time duration. Conversely, in TCT, intervals are differentiated based on their category, with neural dynamics distinctly separating the intervals into two clear categories. Both the parametric representation, akin to a linear attractor, and the categorical representation, characterized by distinct attractor states, are preserved throughout the delay period. Although the circuit represents and sustains intervals using different combinations of neurons, the encoding strategy within each task remains consistent.

Traditionally, the VPC has been primarily recognized for its role in processing and integrating sensory information from various modalities (30, 41, 42, 53), rather than being associated with the neural circuits involved in time estimation (4). Recent evidence has revealed the VPC’s involvement in the encoding of words and phonemes in humans (45), which inherently reflects the importance of this region in the estimation of temporal intervals. Notably, unlike Broca’s area, the VPC has shown promise in predicting speech in patients with conditions like sclerosis that impair communication. This distinction underscores the VPC’s unique ability to manage the temporal dynamics and intonation crucial to speech. Our findings further extend this understanding by suggesting that the VPC is not only capable of integrating multimodal information but also plays a critical role in quantifying time intervals, a function previously unrecognized in non-human primates. To our knowledge, this work provides original evidence that directly links this brain area with the processing of temporal estimates, hinting at a broader functional capacity of the VPC, potentially positioning it as a key player in the neural processing of time-dependent information across different contexts and species.

As discussed in the introduction, previous studies on synaptic connectivity (36–39) and resting-state networks using fMRI (54–56) suggest that the VPC acts as a central hub, receiving inputs from sensory cortices as well as other frontal lobe regions. This strategic positioning allows the VPC to integrate information across multiple sensory modalities, emphasizing its role in complex cognitive processes. However, a key question remains: does the temporal information observed in VPC neurons originate directly from primary and secondary sensory areas, or is it mediated by subcortical regions like the basal ganglia or distant structures such as the cerebellar cortex (57, 58)? The VPC’s ability to process temporal information raises intriguing possibilities about its broader function within neural circuits, especially in relation to time perception and decision-making. Understanding whether the VPC directly receives temporal cues from sensory cortices or whether this information is processed through additional brain structures could significantly advance our comprehension of how time is encoded and utilized in cognitive tasks. Future research should focus on mapping the specific neural pathways that contribute to the temporal coding observed in the VPC. This could involve targeted interventions or tracing studies to delineate the flow of temporal information from sensory inputs through to motor outputs.

Although our results suggest that the VPC, particularly at the population level, has the capacity to maintain time information during working memory intervals, only a limited number of neurons participate in this process (59). These findings imply that the VPC might be more involved in processing and integrating information rather than in the stable and persistent maintenance of temporal codes. While previous studies have shown that the VPC also encodes tactile and auditory sensory information during working memory (30, 41, 42), in our temporal tasks, the number of neurons involved seems even smaller. This observation raises the question of where temporal information is being reliably stored during memory tasks. Previous research highlights two areas as strong candidates for this role: the medial premotor cortex (MPC)(30, 31, 44) and the prefrontal cortex (PFC) (30, 31, 49). For instance, in a context-dependent duration and distance discrimination task (60), the authors found that a subset of PFC neurons exhibited a linear relationship between their firing rates and the presented time intervals. Notably, most of these neurons distinguished between short and long durations in one task context but not in others. Moreover, they identified a relationship between the timing of the first stimulus and the subsequent delay period, suggesting a parametric encoding of time intervals during this retention phase. Although MPC and PFC, have been previously implicated in parametric encoding of physical stimulus features, it remains unclear whether they maintain temporal information in working memory using a strictly linear code at population level. Future studies should investigate the specific roles of the MPC and PFC in time encoding memory, potentially revealing how these regions interact with the VPC to support temporal processing.

Additionally, interactions between cortical areas could be explored by recording the oscillatory activity of local field potentials (LFPs) during TICT and TCT. Recordings in the MPC and PFC suggest that the beta band (15 to 30 Hz) in the VPC might encode both categorical and parametric multimodal information (61, 62). However, the contribution of the alpha band (8 to 14 Hz) to this process remains unclear. Furthermore, we propose that examining the interaction between different brain regions through LFP-spike coherence (62, 63) could yield valuable insights into how distinct neural circuits contribute to time encoding. Exploring how these oscillatory patterns influence neuronal firing and interact across regions could provide a deeper understanding of the specific roles various cortical areas play in processing and maintaining time information in TICT and TCT.

We would like to emphasize the pivotal role of recurrent neural networks (RNNs) in advancing our understanding of temporal information processing (11, 15, 64–67). Previous RNN simulations of analogous tasks (64, 68) have revealed a similar separation between linear and categorical dynamics, mirroring the geometric patterns observed in our experimental results. This suggests that comparable dynamics emerge in both RNNs and neuronal populations across tasks. However, a crucial question remains: how do neural circuits transition from linear and parametric storage to categorical representations of information? Recent studies have provided insights into how RNNs adapt across different tasks (66, 69). Although further experiments and geometric analyses are needed to fully unravel these transitions in biological networks (70–72), RNNs provide a powerful framework for exploring the flexibility of neural circuits across various contexts (69).

In summary, the VPC’s population activity encodes time information, displaying distinct neural dynamics based on task demands. In TICT, the VPC maintains a parametric representation, continuously encoding temporal intervals. In contrast, during TCT, the VPC preserves categorical distinctions, segmenting time intervals into discrete categories. These findings suggest that the VPC flexibly adapts its processing to meet cognitive requirements, potentially integrating temporal information from multiple sources. Intriguingly, this adaptive neural flexibility mirrors Einstein’s concept of time dilation, where the perception of time varies relative to the observer’s frame of reference. Just as Einstein revolutionized our understanding of time as not absolute but relative and context-dependent, our findings indicate that the brain’s temporal encoding is dynamic and shaped by cognitive context. The VPC’s ability to shift between parametric and categorical representations underscores the brain’s remarkable capacity to adapt to perceptual time demands.

## ACKNOWLEDGMENTS

This work was supported by grants PAPIIT-IN210819 (to R.R. and R.R.-P.), PAPIIT-IN205022 (to R.R.-P.) and IN203825 (to R.R.-P.) from the Dirección de Asuntos del Personal Académico de la Universidad Nacional Autónoma de México and CONAHCYT-240892 (to R.R.), CONAHCYT-319347 (to R.R.-P.) from Consejo Nacional de Ciencia y Tecnología; IBRO Early Career Award 2022 (to R.R.-P.) from International Brain Research Association. H.D. is a doctoral student from Programa de Doctorado en Ciencias Biomédicas, UNAM. L.B. is a postdoctoral student (Postdoctoral fellowship CONACYT-838783).

## MATERIALS AND METHODS

### Time Interval Comparison Task (TICT)

In the TICT, two monkeys (Macaca mulatta) were trained to compare the durations of two successive intervals separated by a fixed delay period of two seconds. The setup involved the monkey seated in a primate chair, with its right arm, hand, and fingers securely restrained. Its left hand was used to operate an immovable key, positioned at about a 90-degree angle at the elbow, along with two push-buttons spaced 3.5 cm apart from center to center and located 18 cm away from the key, directly in front of the animal at eye level. Stimulation was delivered through a computer-controlled mechanical stimulator, which moved a probe with a 2mm rounded tip across the skin of a restrained finger, applying a consistent pressure of 150 µm. The trial began when the probe contacted the skin (“PD”), prompting the monkey to initiate the task by placing its non-stimulated hand on a touch-sensitive lever (“KD”). After a random waiting period of 1.5 to 3.0 seconds, which prevented the monkey from anticipating the stimuli, two consecutive pulses, each lasting 20 ms, were delivered to the indented skin. The time interval (*int1*) was defined by the separation between these two pulses. Each of these intervals lasted between 400 and 2000 milliseconds, with each individual pulse spanning 20 milliseconds. The monkeys was required to compare these intervals (*int1* and *int2*), which were interspersed by a 2-second working memory period. Following the comparative phase, there was an additional 1-second delay in which the monkeys had to maintain its decision until the probe up event (“PU”). This event signaled the “GO” cue, prompting the monkeys to release the immovable key (“KU”) and indicate its decision by pressing one of two push-buttons. The choice depended on whether the monkeys perceived the second interval as longer (*int1*<*int2*, lateral switch) or shorter (*int1*>*int2*, medial switch) than the first. Correct decisions were rewarded with drops of apple juice.

### Time Categorization Task (TCT)

The TCT has been described before (73). Monkeys were trained to categorize the duration of a vibrotactile stimulus as either “Long” or “Short”. The monkey underwent trials arranged in blocks, termed “sets,” which consisted of predetermined groups of stimuli. Before each set commenced, the monkey was shown the shortest and longest stimuli to establish the range for temporal categorization. The training employed two different ranges, which remained constant within a set. Vibrotactile stimuli consisted of short mechanical pulses, each a 20ms single-cycle sinusoid at 20 Hz, lasting from 50 ms to 550 ms in the short temporal range (SRT) and from 330 ms to 1170 ms in the long temporal range (LRT). Each trial initiated with a probe down event (“PD”), where the probe indented the skin to a depth of 150 µm. This was followed by a randomly timed waiting period (1.5-3s) to prevent anticipatory responses from the monkey. After this, the intervals for categorization were delivered, followed by a fixed 2s delay (Fig. 1B). The completion of this delay was indicated by the probe up event (“PU”), which acted as the “GO” cue, prompting the monkeys to release the immovable key (“KU”) and make its choice using one of the two push-buttons. Correct responses were rewarded with drops of fruit juice.

In both tasks, TICT and TCT, animal’s handling was consistent with the standards of the National Institutes of Health and the Society for Neuroscience, and all protocols were approved by the Institutional Animal Care and Use Committee of the Instituto de Fisiología Celular at the National Autonomous University of Mexico (UNAM).

### Recordings

Neuronal recordings were obtained using an array of seven independently movable microelectrodes (2–3 MΩ) inserted into the ventral premotor cortex (VPC)(36–38, 41) (Fig. 1E) in both the contralateral (right hemisphere) and ipsilateral (left hemisphere) sides relative to the stimulated hand in monkeys RR22 and RR34. The exact placement of the electrodes was confirmed through standard histological techniques. Recording sites varied across sessions, with each term consisting of approximately 30 sessions (1 session per day). A total of 45 recording sessions were conducted for monkey RR22 and 68 sessions for monkey RR34. The locations of electrode penetrations were used to construct surface maps of each cortical area, achieved by marking the edges of the small chamber (7 mm in diameter) placed over each cortical region. In total, we recorded 180 neurons during the TCT task and 702 during the TICT task, including data from both monkeys, RR22 and RR34. It is important to note that neurons encoding interval information were distributed throughout the VPC, making the identification of specific subgroups of responsive neurons unlikely.

### Data Analysis

For the analyses described below and the results presented in this study, only hit trials were considered. During the categorization task, trials corresponding to *int=*300 ms for SRT and *int*=750 ms for LRT were separated based on the animal’s decision.

#### Firing Rate

For each neuron, a time-dependent firing rate was calculated for each trial using overlapping rectangular causal windows with a length of 200 ms and steps of 20 ms. For visualization purposes only, in Figs. 2, 3, S1, and S2, a causal Gaussian kernel with a σ of 200 ms was applied.

#### Mutual Information

For each neuron, we calculated the relationship between the firing rate (*r*) and the interval duration (*int*) for each window using Shannon mutual information (74, 75). This measure allows us to study the nonlinear relationships between both variables. In the raster of all neurons, we indicate the time bins where the different neurons carry significant temporal information. In Figures 4 and S3, we compute the percentage of neurons with significant interval-associated information (*I_Int_*). To calculate *I_Int_*, we considered the firing rate probability distributions associated with the different intervals (*P(r|int)*) and the global firing rate probability distribution (*P(r)*) by combining trials from all intervals values:

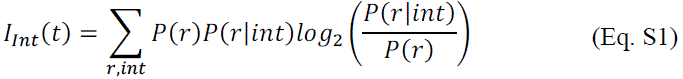

The significance of mutual information was calculated per neuron in each time bin. A permuted value of mutual information was obtained by performing 1000 permutations of the trials. A mutual information value was considered significant if the probability of a permutation having the same or greater value was below 0.05 (p < 0.05). Additionally, to address the issue of finite sampling, we applied the correction proposed in (74). For the significance test, we also performed a correction for multiple comparisons using a clustering method previously described (76). This was done by retaining only the set of significance-connected time bins with a size larger than a defined threshold.

#### Coding with F-test

In Figures 4A, 4B and S3A, and in the various neurons shown in the raster plots (Figures 2, 3, S1, and S2), significant coding for time intervals in the firing rate was also computed using the F-test metric (indicated as “Int Cod” in orange). The F-test allows us to compare the firing rate (*r*) variance within different data groups, corresponding to the different intervals (*int*), with the variance of the mean firing rates for each interval. To determine significance, a permutation test with *p<0.05* was applied.

#### Population States Similarity

To analyze the similarity of population states across different times of the task (Fig. 4B, 4D and S3B), we computed, for each time window of the delay in TICT and TCT, a population vector with the form:

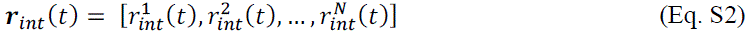

where each element of this vector corresponds to the mean firing rate of all hit trials of each neuron i for interval *int*, at that specific time 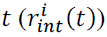. To compare the population state across all intervals, we concatenated the response associated with each interval (*int*) for every neuron i at each time point. This results in a vector with dimensions *N* × *c*, where *N* is the number of neurons and *c* is the number of intervals (*int*). The comparison between population states was assessed by computing the cosine similarity metric. To measure the similarity between two vectors, the cosine of the angle between them can be used. This measure calculates the cosine of the angle between two vectors, without considering their magnitudes. The cosine similarity (*CS*) is 1 for two parallel vectors and 0 for orthogonal vectors (90° difference). The cosine similarity can be mathematically represented as:

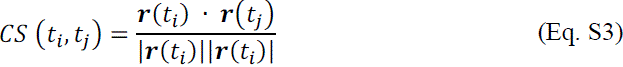

where *r*(*t*_i_) · *r*(*t*_j_) is the dot product of two vectors at times *t*_i_ and *t*_j_ respectively, and |*r*(*t*_i_)| and |*r*(*t*_j_)| are the magnitudes of the vectors at those same time points. Therefore, *CS* (*t*_i_, *t*_j_) defines a matrix of dimension *T* × *T*, which allows for quantifying the similarity of population states at two different times.

#### Principal Component Analysis (PCA)

The main goal of applying PCA to electrophysiological data during cognitive tasks is to find a new coordinate system in which the data can be represented more concisely and compactly. In other words, the aim is to define a low-dimensional subspace that captures most of the variance from the high-dimensional neural state space. Typically, the number of significant dimensions is reduced substantially, going from as many possible dimensions as neurons (*N*) to just a few axes that concentrate most of the variance (51). To characterize how the population activity varies for different interval values (*int*) as a function of time (*t*), we applied PCA (Figures 5, 6, and S4). PCA produces a new coordinate system for the N-dimensional data, where the first coordinate explains the largest portion of the variance in the neural population. The second coordinate accounts for the next largest portion of the variance, and so on; however, each subsequent axis is constrained to be orthogonal to all the previous ones. PCA was computed from the covariance matrix of the firing rate, which averages over time windows (*t*) and intervals (*int*):

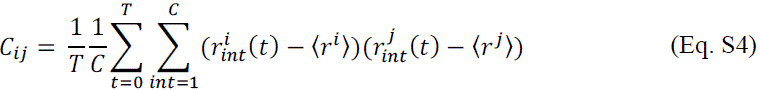

where *T* is the number of time windows in the period being considered, *C* is the number of different intervals recorded in each experiment (7 in TICT, 12 in SRT, 8 in LRT). *r*^i^ (*t*) is the average firing rate of neuron i for the interval *int* at time *t*, and 〈*r*^i^〉 is the average firing rate of neuron i across all intervals and time windows. The diagonalization of the covariance matrix, *C* = *UDU*^*T*^, produces a new coordinate system given by the columns of matrix *U*, which we call the derived axes or principal components (PCs). On the other hand, *D* is a diagonal matrix of positive values. The diagonal elements of *D* represent the amount of variance in the population activity captured by the corresponding PCs. We then order the PCs according to the amount of variance captured. The projection of the N-dimensional data onto the *k*-th PC is given by:

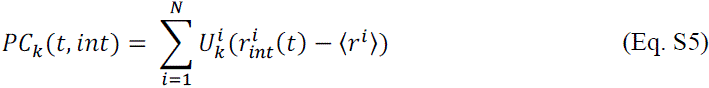

where 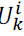 is the i element of the *k* PC (*U*_*k*_). Therefore, the PCs are linear readouts of the population activity; in other words, they are linear combinations of the firing rates of the individual neurons. Thus, the contribution of each neuron to a given *PC*_*k*_ is represented by the i-th element of *U*_*k*_. These PCs can be viewed as a low-dimensional description of the population activity in this coding subspace.

In both tasks, TICT and TCT, we divided the population study into two parts. In the first stage, we focused on the relevant dynamics during the presentation of the intervals (Figure 5 and S4A). We computed the covariance matrix (Eq. S4) using the neuronal activity from the beginning of the interval (*int*) to 200ms after its end. In other words, we only summed over the time windows that included the presented intervals (*int*). It is necessary to include these additional 200ms since we use deterministic windows and the response latency of the VPC is greater than 150ms (31). Next, we focused on the study of the working memory delay period in TICT and the decision maintenance period in TCT (Figure 5 and S4A). I In this case, we computed the covariance matrix (Eq. S4) using the time windows from these periods shifted 200 ms forward in the timeline to compensate for VPC’s response latency.

#### Noise and Dimensionality

To isolate the principal components (PCs) that capture significant fluctuations in neural activity, we first determined a ‘noisy’ firing rate for each neuron. This was achieved by calculating the firing rate of each neuron at time *t* during a trial within the interval *int*, and subtracting the average firing rate of all trials at the same time point *t* for that interval *int*. This process removed the signal component relevant to the neuron’s encoding function across intervals (*int*) and time, leaving only the residual fluctuations. Then, for each neuron, we randomly selected twenty trials to compute the covariance with every other neuron. This random selection was repeated in each iteration to ensure a robust estimation of each neuron’s contribution to the noise covariance matrix, quantifying the covariance across all possible neuron pairs. Mathematically, this follows:

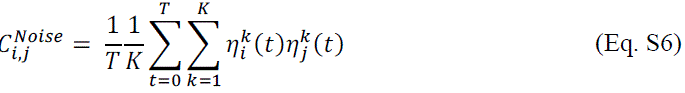

where 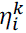 is the noisy firing rate value in time *t* of trial *k* that belongs to neuron i. PCA was then performed on the resulting covariance matrix to extract the eigenvalues, which indicate the variance captured by the principal components (PCs). We calculated the proportion of explained variance for each derived PC. To ensure statistical robustness, this matrix construction and eigenvalue extraction process was repeated 100 times with varying random seeds, allowing us to determine the mean and standard deviation of the noise-associated explained variance for the different PCs. This analysis facilitated the determination of the system’s dimensionality by identifying the principal components (PCs) with eigenvalues exceeding the noise threshold, which were deemed statistically significant. These significant PCs were subsequently used to compute the relevant projections.

#### Speed and Distance

To calculate the speed and distance of the neural trajectories, we utilized the Euclidean distance within the subspace formed by the first significant principal components (PCs). For the absolute metrics, we directly compared the trajectories at the same time points *t*_i_, measuring differences across different values of intervals (*int*) in the neural activity evolution over time in the reduced dimensional space. This approach allowed us to capture the differences in neural population dynamics based on the temporal progression of interval-related activity. Based on these differences, the speed shown in Figure 5A, B and S4A was calculated.

The distance between the final positions of the population vector at the end of each interval, as shown in Figures 5A, B, and S4A, was calculated (15). For each task, an intermediate interval was used as the reference: 1000 ms for TICT, 300 ms for SRT, and 750 ms for LRT. First, a projection axis was determined using the difference vector between the reference vector and the vector corresponding to 1300 ms for TICT, and for SRT and LRT, the vector assigned to the different category from the reference vector. Next, the difference vector between the final position of the reference vector and the final positions of the other intervals was computed. To obtain the signed distance, we took the dot product of these vectors with the normalized vector of the projection axis. It is important to note that in these figures, the distance for the population vector of the reference interval is 0.

For the relative metrics, we employed the Kinematic Analysis of Neural Trajectories (KiNeT) as described in reference (15). The KiNeT method aligning neural trajectories based on their dynamics rather than strictly on external time cues. This is particularly important when comparing neural responses to stimuli of different durations or when assessing how neural representations evolve during tasks requiring temporal judgments. First, for each trajectory j, we computed the time point where it exhibited the greatest similarity to a reference trajectory. This was achieved by finding the time 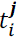 that minimized the Euclidean distance between the two trajectories in the principal component subspace. By comparing the time of maximum similarity 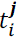 with the absolute time *t*, we derived a measure of the trajectory’s speed. If the trajectory reached the point of maximum similarity earlier than the reference trajectory, it was considered faster, and vice versa. Using this calculation, the speed in Figures 6A, B and S4B was computed.

To calculate the signed distance, we measured the Euclidean distance between the trajectories at this relative time. We then assigned a sign to the distance based on the dot product between the difference vectors of the compared trajectories and the difference between the reference trajectory and the first trajectory. This approach enabled us to capture not only the magnitude of the differences between trajectories but also their directional relationship, indicating the direction in which they were moving relative to one another. We used this calculation in Figures 6A, B and S4B.

---

**Figure S1.**
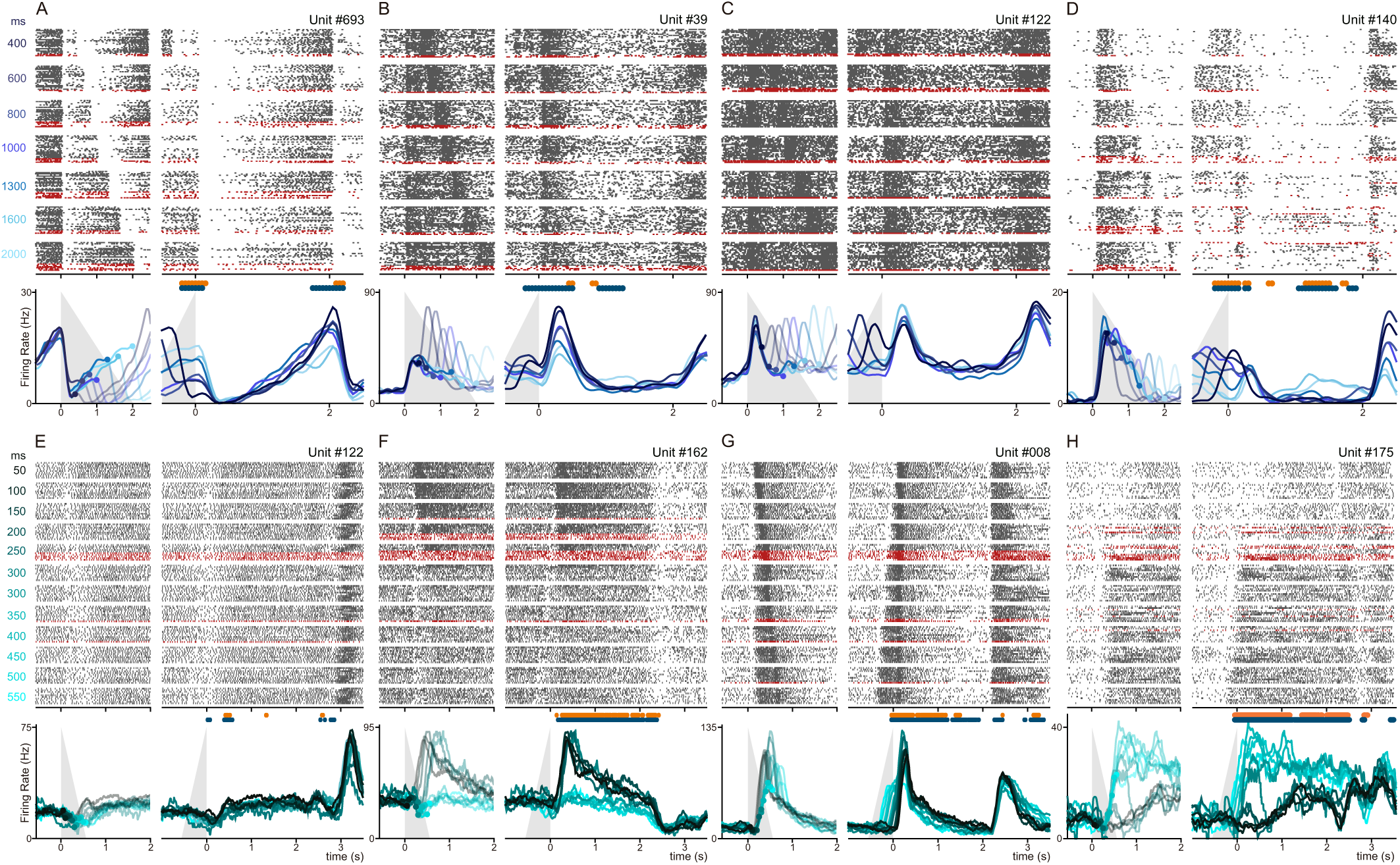
VPC single neurons’ activity during TICT and TCT. Raster plots and corresponding firing rate profiles of eight individual neurons, with four (A-D) during TICT and four (E-H) during SRT. Black and red ticks represent spikes during trials with correct and incorrect answers, respectively. In each neuron, left raster plots denote activity aligned to the beginning (t=0 s) of the interval (*int1* or *int*) presented, while right rasters aligned activity to the start of the fixed 2-second delay. Firing rate plots beneath each raster was computed as the mean of correct-answer trials for each interval value used. Grey triangles indicate the crescent interval values, visually representing how firing rates vary with changes in interval duration across successful trials. Orange and green markers over activity profiles denote time-windows with significant (p<0.05) mutual information and F-test. Note how neurons from A to D during TICT are involved in the encoding of time intervals. For example, neuron A increases its firing rate as the interval lengthens, while unit B fires at a higher rate as the interval decrease. Neurons C and D also mark the interval, albeit with some processing variations. In the STR context, F and H neurons are involved in categorizing the time intervals and carrying the category decision throughout of the delay. On the other hand, neuron E shows its peak activity during the execution of the decision after the decision-delay period, and neuron G performs more complex computations, arguably related with the encoding of the time interval’s identity. This shows once more the wide heterogeneity of responses in VPC during TICT and TCT.

**Figure S2.**
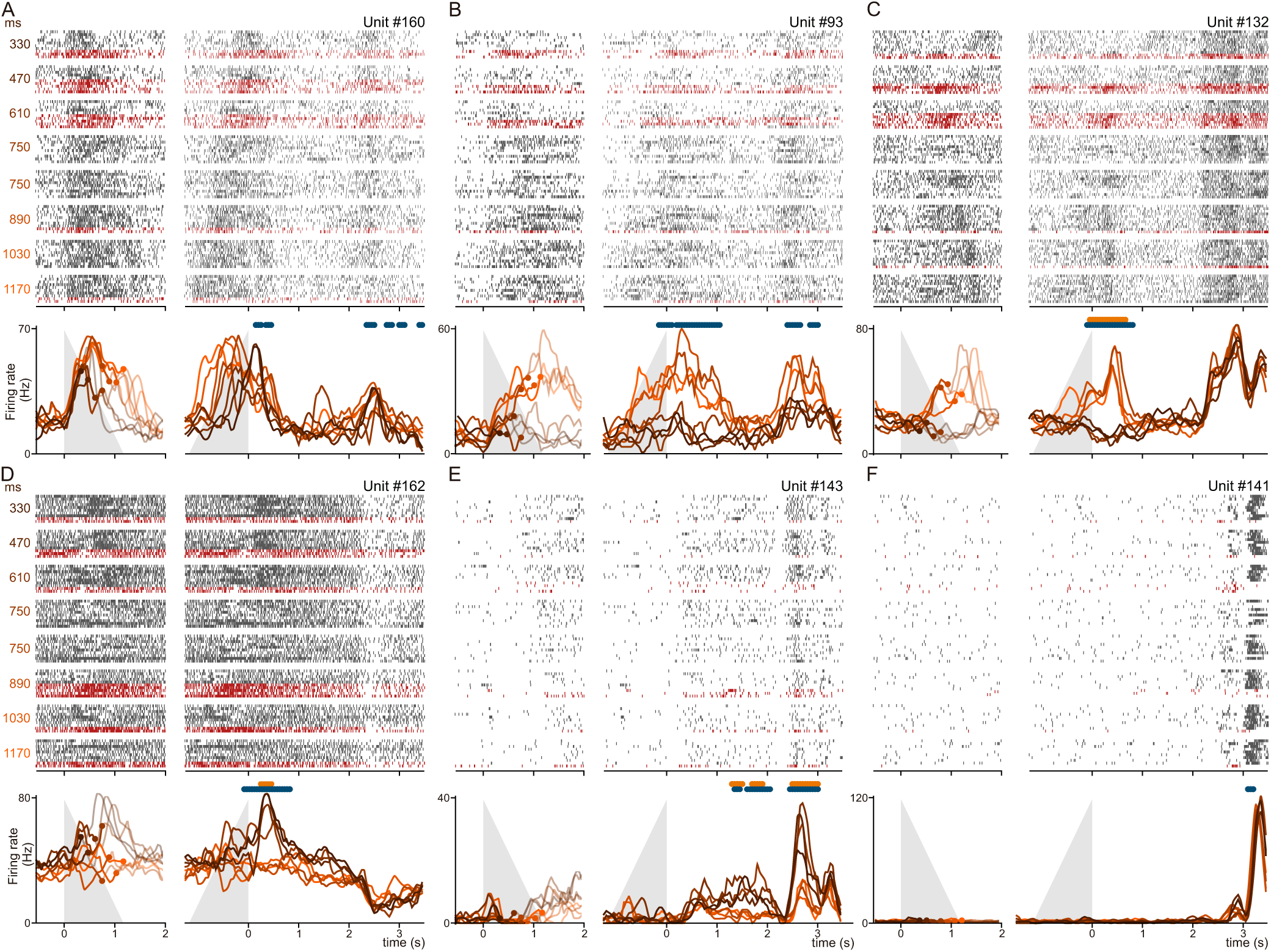
Individual neurons from the VPC during the LTR in TCT. (A-F) Raster plots and corresponding activity profiles of six individual neurons during the long temporal range (LTR) in the TCT. Black and red ticks represent spikes during trials with correct and incorrect responses, respectively. For each neuron, left raster plots denote activity aligned at the beginning (t=0 s) of the interval (*int*), while in right rasters activity was aligned to the start of the fixed 2-second decision delay. Trials are organized by the duration of *int*, with durations labeled on the left side (ranging from 330 to 1170ms). In this set, all intervals exceeding 750ms should be categorized as long. The 750ms trials, which serve as the thresholds, were divided based on the animal’s decision. Firing rate plots beneath each raster were computed as the mean of correct-answer trials for each of the twelve interval values used. Grey triangles indicate the crescent interval values, showing how firing rates vary with changes in interval duration among successful trials. Orange and green markers on the neuronal activity denote time-windows with significant (p<0.05) mutual information and F-test. Neurons B, C, and D encode the category of the interval, particularly during the first part of the decision delay. Neuron A exhibits dynamics more involved in quantifying the duration of the intervals, as evidenced by the detailed activity in the raster plots. Neurons E and F primarily respond after the probe up (PU) event and would be involved in the execution of the decision.

**Figure S3.**
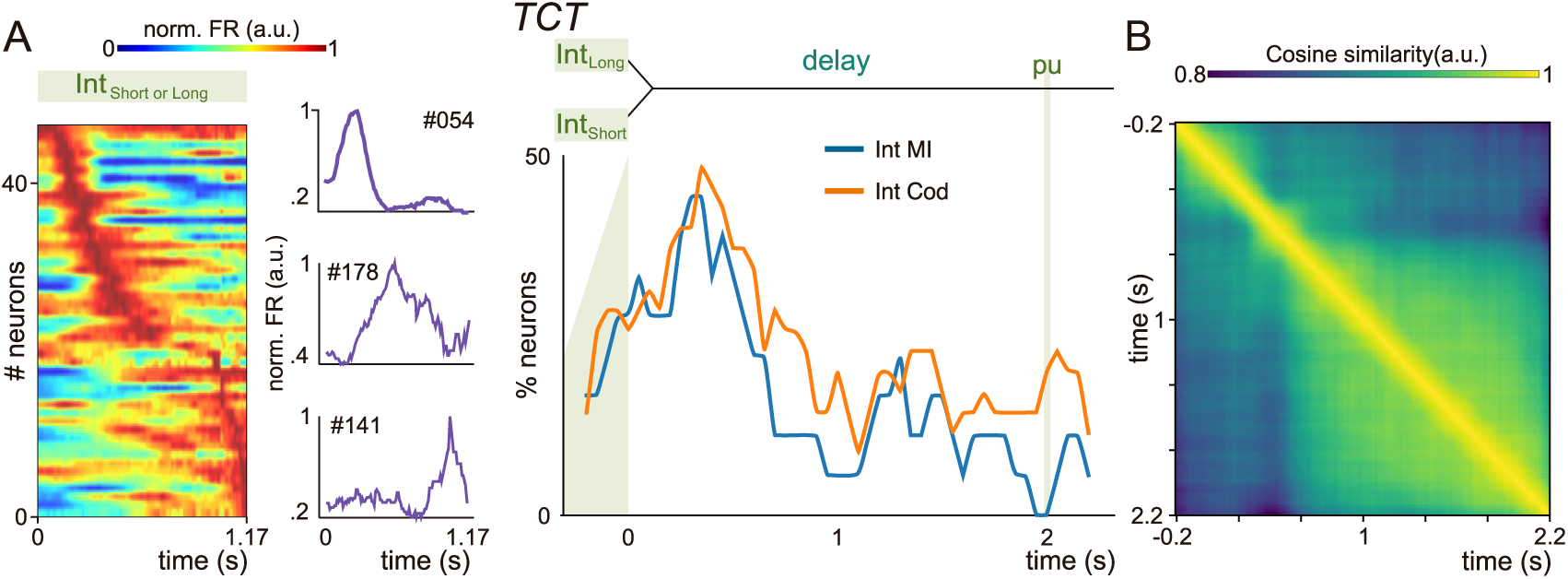
Estimation and maintainance of temporal information by the population of neurons in the LTR context. (A) **Left:** Dynamics of individual neurons during the presentation of the interval to be categorized in the LRT (TCT). Neurons with more than 5 from the basal activity were normalized and positioned on the color-coded graph by the time of peak activity. Neurons at the top respond earlier, while those at the bottom respond later. A “moving bump” dynamic is used by the population of neurons as a time-interval encoding strategy. **Middle**: Example of neurons with normalized peak activity at three different specific times within the range of intervals presented. **Right**: percentage of neurons with significant coding during the decision maintenance delay of the LRT. Mutual information (blue) and F-test (orange) were used to identify neurons with significant time-interval encoding (p<0.05). Consistent with Figure 4, a large number of neurons demonstrate significant encoding at the start of the delay, with a gradual decline over the 2-second period. However, a substantial level of significant encoding remains throughout the delay. (B) Cosine similarity: A population vector is generated at each time point, with each line reflecting a neuron’s activity for different intervals (*int*). The cosine of the angle between vectors from two different times is visualized, with yellow indicating high similarity and blue showing low similarity among population states. As noted in Figure 4, there is a marked separation between the population state at the end of the interval and the beginning of the delay, with respect to the dynamics throughout the subsequent maintenance periods.

**Figure S4.**
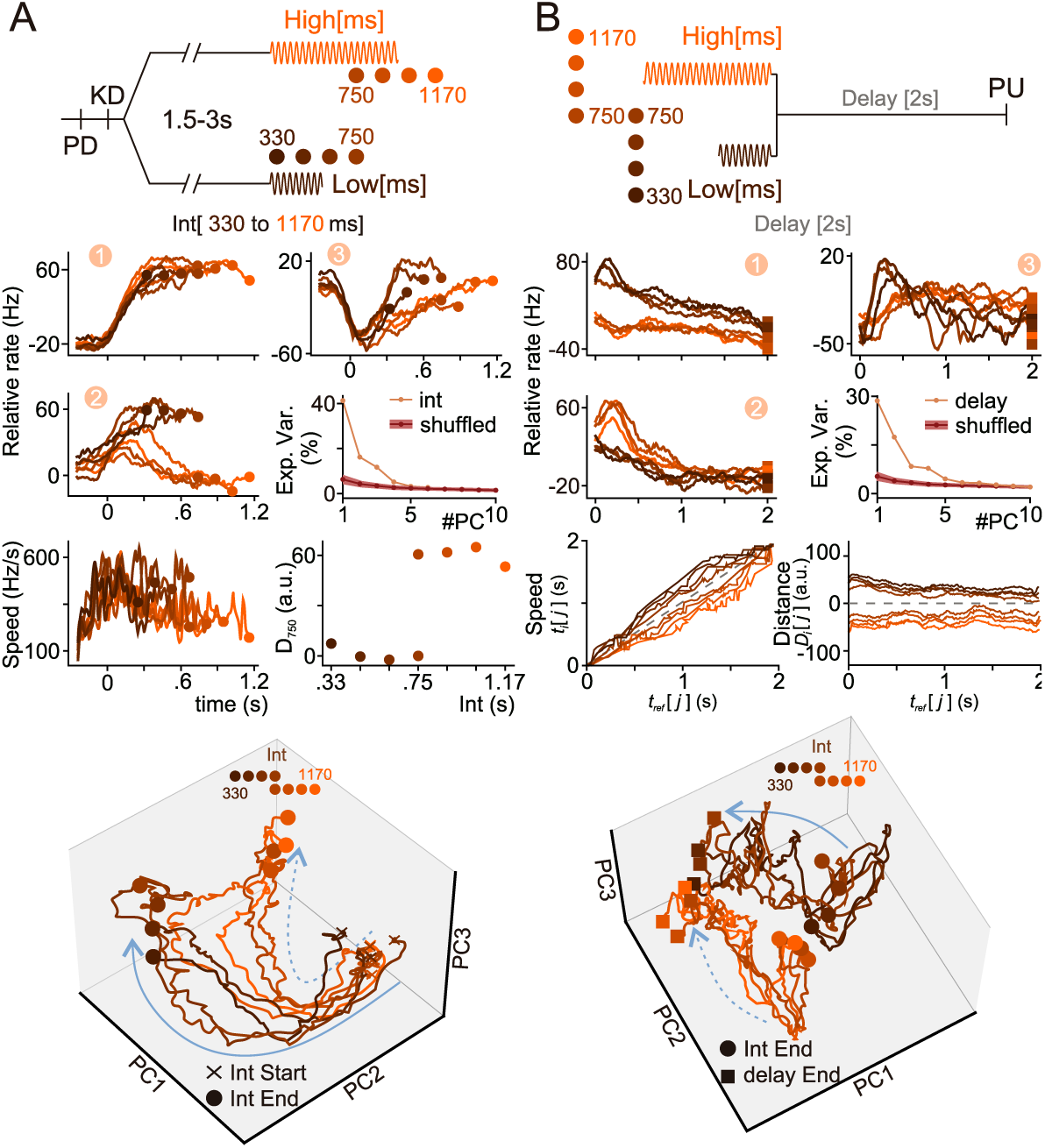
Dynamics involved in estimation and maintenance of time-interval information during LRT. Population dynamics of the VPC during the presentation of time intervals (A) and in the subsequent decision maintenance delay (B) for the LRT context. Principal components (PCs) were derived from applying PCA to the covariance matrices constructed using only the activity from the intervals presented plus 200 ms from the memory delay (panel A) and from the entire 2-second decision maintenance delay, shifted 200 ms forward in the time line. PCs were ordered according to their EV for both, A (PC1:41.3%, PC2: 16.2%, PC3:11.7%) and B (PC1: 28.6%, PC2: 17.4%, PC3: 8.4%) panels. **Top:** Schematic representation of the period being studied: interval representation (A) or decision maintenance delay (B). **Middle:** Individual projections of neural activity framed in: each interval (A) and decision-maintenance delay (B), into the three significant PCs. Note how a categorical representation emerges at the end of PCs 2 and 3 in panel A, and this representation is maintained throughout the entire delay in PCs 1 and 2 in panel B. **Bottom:** 3D-phase diagram using the projections in the three significant dimensions traces the evolution of neural dynamics during the interval period (A) and delay (B) of the LRT. (A) Dynamics start in the same regions (crosses) and ends finish (circles) in different positions of the state space depending on the interval category. The distance from the final position of the 750ms interval (D_750_) reveals a clear categorical separation of the intervals. (B) Dynamics in the delay begin (circles) at different positions, determined by the endpoint of the interval period shown in panel A, and end (squares) in the same regions of the space state, denoting the proximal execution of the decision. While the speed remains approximately constant for both categories throughout the delay, a marked distance between them is maintained, emphasizing distinct neuronal representations for each category.

